# Spatial tumour gene signature discriminates neoplastic from non-neoplastic compartments in colon cancer: unravelling predictive biomarkers for relapse

**DOI:** 10.1101/2022.09.27.509641

**Authors:** Katja Sallinger, Michael Gruber, Christin-Therese Müller, Lilli Bonstingl, Elisabeth Pritz, Karin Pankratz, Armin Gerger, Maria Anna Smolle, Ariane Aigelsreiter, Olga Surova, Jessica Svedlund, Mats Nilsson, Thomas Kroneis, Amin El-Heliebi

**Affiliations:** Division of Cell Biology, Histology and Embryology, Gottfried Schatz Research Centre, Medical University of Graz, Graz, Austria; Center for Biomarker Research in Medicine (CBmed), Graz, Austria; Division of Oncology, Department of Internal Medicine, Medical University of Graz, Graz, Austria; Department of Orthopaedics and Trauma, Medical University of Graz, Graz, Austria; Diagnostic and Research Institute of Pathology, Medical University of Graz, Graz, Austria; Science for Life Laboratory, Department of Biochemistry and Biophysics, Stockholm University, 17165, Solna, Sweden; 10x Genomics, Life City, Solnavägen 3H, 113 63 Stockholm, Sweden; Biotechmed Graz, Austria

**Keywords:** *in situ* sequencing, spatial transcriptomics, colon cancer, predictive biomarker, tumour compartment, tumour gene signature

## Abstract

**Background:** Therapeutic management of stage II colon cancer remains difficult regarding the decision whether adjuvant chemotherapy should be administered or not. Low rates of recurrence are opposed to chemotherapy induced toxicity and current clinical features are limited in predicting disease relapse. Predictive biomarkers are urgently needed and we hypothesise that the spatial tissue composition of relapsed and non-relapsed colon cancer stage II patients reveals relevant biomarkers.

**Methods:** The spatial tissue composition of stage II colon cancer patients was examined by i*n situ* sequencing technology with sub-cellular resolution. A panel of 175 genes was designed investigating specific cancer-associated processes and components of the tumour microenvironment. We identified a tumour gene signature to subclassify tissue into neoplastic and non-neoplastic tissue compartments based on spatial expression patterns generated by *in situ* sequencing (GTC-tool – Genes-To-Count).

**Results:** The GTC-tool automatically identified tissue compartments that were used to quantify gene expression of biological processes upregulated within the neoplastic tissue in comparison to non-neoplastic tissue and within relapsed versus non-relapsed stage II colon patients. Three differentially expressed genes (*FGFR2, MMP11* and *OTOP2*) in the neoplastic tissue compartments of relapsed patients in comparison to non-relapsed patients were identified predicting recurrence in stage II colon cancer.

**Conclusions:** In depth spatial *in situ* sequencing revealed novel potential predictive biomarkers for disease relapse in colon cancer stage II patients. Our developed open-access GTC-tool allows to accurately capture the tumour compartment and quantify spatial gene expression in colon cancer tissue.

## Introduction

Colorectal cancer is the third most commonly diagnosed cancer, and the second leading cause of cancer associated mortality worldwide [1]. The incidence of colorectal cancer is increasing each year by a factor of 0.4% in the European Union, which is associated to increased life style effects [2]. Overall survival is strongly associated with stage at diagnosis, estimated at 13%, 31%,32% and 24% for stages I, II, III and IV, respectively [2]. Therapeutic management has significantly improved over the last decade, including advances in screening, (neo) adjuvant treatment, targeted- and immune checkpoint therapies [3]. Despite these achievements, treatment decision remains difficult in colon cancer stage II, representing roughly 1/3 of all cases, especially with regards to adjuvant chemotherapy [4]. Overall, low rates of tumour recurrence are opposed to potentially toxic effects of chemotherapy. Adjuvant therapy for all patients with diagnosed stage II colon cancer would be an overtreatment, leading to reduced quality of life, little measurable benefit in outcome and unnecessary exposure to chemotherapy-induced toxicities [5]. Hence, current guidelines rely on high-risk clinic-pathological features to identify patients who would more likely benefit from adjuvant treatment [6]. However, high-risk features, such as T4 tumours, perineural or lymphovascular invasion, poorly or undifferentiated tumour grade, intestinal obstruction or tumour perforation are unreliable in predicting benefit of adjuvant treatment [7]. Vice versa, low-risk patients may develop tumour recurrence quite early after surgery [7]. Therefore, more precise biomarkers indicative for early recurrence in stage II colon cancer are needed [6]. Various biomarker panels have been examined for the purpose of recurrence prediction such as immune scores [8], 12-gene recurrence score assays [9] or 18-gene expression based classifier ColoPrint [10]. These approaches are based on either small antibody combinations (CD3 + CD8 immunohistochemistry), or bulk tissue analysis using reverse transcription–polymerase chain reaction (RT-PCR) or array based approaches [6,11]. However bulk analysis only informs about the average sub-clonal composition, with strong bias towards the largest clones present [12]. Similarly, the spatial histological architecture is lost in bulk RNA or DNA analysis due to tissue lysis [13]. Thereby important biological processes, such as proliferation or metabolic dysregulations in cancer are only measurable indirectly without their spatial interactions [14]. However, understanding the spatial expression patterns of neoplastic tumour tissue and their surrounding microenvironment on a subcellular level can improve the knowledge of disease recurrence [13, 14]. Our knowledge of the basic molecular mechanisms in colon cancer has increased substantially within the last years and was revolutionized by technologies such as genome-wide expression analysis and single cell RNA sequencing (scRNAseq) unravelling the complexity of colon cancer [15, 16]. Using scRNAseq, the transcriptomic diversity of different cell types within a specimen has been revealed in high detail but a major drawback remains the dissociation of tissue structure, again losing the spatial context. To overcome this issue, spatial transcriptomics approaches have been developed such as hybridisation-based *in situ* sequencing (HybISS) and direct RNA hybridisation-based *in situ* sequencing (dRNA-HybISS) allowing for highly multiplexed spatial mapping of transcripts within tissues [17,18].

Here, we hypothesise that the spatial histological expression patterns of relapsed and non-relapsed colon cancer stage II patients are different. Therefore, we aimed to investigate the spatial tissue composition in yet unprecedented resolution by dRNA-HybISS down to single-cells and –molecules, beyond current spatial transcriptomics approaches in colon cancer stage II patients. First we designed a panel of 175 genes to examine the expression of various important biological processes in colon cancer including angiogenesis, apoptosis, proliferation, stemness, hypoxia, oxidative stress, invasion, and energy metabolism as well as markers for components of the microenvironment including cancer associated fibroblasts. Based on the expression pattern of the targeted ISS genes, in particular a tumour gene signature, tissue compartments are automatically generated to subclassify the investigated tissue into neoplastic and non-neoplastic compartments. By using these gene expression-based tissue compartments, we are able to quantify gene expression related to biological processes upregulated within the neoplastic tissue in comparison to non-neoplastic tissue. Second, we statistically evaluated which spatially differential expressed genes are predictive for tumour recurrence.

Summarized, we identified a spatial tumour gene signature and developed a computational tool to classify tissue into neoplastic and non-neoplastic tissue by *in situ* sequencing informed expression. We thereby identified *FGFR2, MMP11* and *OTOP2* as three differentially expressed genes in the neoplastic tissue predicting tumour recurrence in stage II colon cancer.

## Materials and Methods

### Study design – Patient cohort

For this retrospective study, 10 patients were selected with diagnosed stage II colon cancer from the Biobank Graz. Each patient was observed for at least 3 years after tumour resection and their final tumour recurrence status labelled as either relapsed (5 patients, 50%) or non-relapsed (5 patients, 50%). None of the patients received adjuvant chemotherapy after tumour resection. Tumour tissue was formalin fixed and paraffin embedded (FFPE) and stored in the Biobank Graz. Neoplastic and non-neoplastic colonic tissue was present in each tissue section.

### Ethic approval

The study protocol was approved by the Ethics Committee of the Medical University of Graz (29-187 ex 16/17) and written informed consent was obtained by all patients. This study follows the ethical principles as outlined in the declaration of Helsinki and good clinical practice.

### Panel design

A panel of padlocks was designed to target 175 transcripts, intended to investigate different biological processes within the tumour and its microenvironment. It includes genes involved in angiogenesis, apoptosis, autophagy, necrosis, proliferation, oxidative stress, hypoxia, stemness, invasion, epithelial-mesenchymal transition (EMT), energy metabolism as well as different epithelial cells, tumour associated stromal cells and immune cells (Supplementary Table 1). The exact target sequences and padlock probes design is propriety information by Cartana (Stockholm, Sweden, now part of 10x Genomics, California, USA) and are not known by the authors.

### Tissue preparation and ISS Library preparation

For application of the *in situ* sequencing method 5μm FFPE tissue sections were processed according to the manufacturer’s protocols (Cartana, part of 10x Genomics). All reagents were provided in kits (High Sensitivity library preperation kit) by Cartana. In short, sections were baked at 60°C for one hour, deparaffinised in xylene or Histolab Clear (Sanova Pharma, Vienna, Austria), rehydrated in 100%, 85% and 70% ethanol and permeabilised in a steamer by using citrate buffer of pH 6 for 45 minutes. The sections were then dehydrated in an ethanol series and air-dried in order to attach hybridization chambers onto the slides and to form sealable reaction chambers (Secure Seal, Grace Biolabs, Oregon, USA). All hybridisation steps were performed in RNAse free humidity chambers. Padlock probes were then directly hybridised to the RNA at 37°C overnight, followed by ligation at 37°C for 2 hours. After the ligation process, a circular oligonucleotide was formed and amplified overnight at 30°C in a rolling circle amplification (RCA) reaction, resulting in RCA products (RCP).

### Sequencing and stripping

All used reagents were provided by Cartana in a kit (Sequencing kit). Adapter probes were hybridised at 37°C for 1 hour, followed by a washing step with washing buffer 2. Afterwards the sequencing probes were hybridised at 37°C for 30 minutes. The sections were washed with washing buffer 2,mounted with SlowFade Gold Antifade Mountant (Thermo Fisher Scientific, Massachusetts, USA) and imaged. The procedure for every sequencing cycle is as follows: after each sequencing cycle, adapter- and sequencing-probes were stripped off by adding three times 100% formamide to each slide for 1 minute at room temperature. This measure was followed by a washing step with washing buffer 2. The ISS cycles were in total repeated 5 times, with 5 different adapter probe pools and imaged in 5 channels (DAPI, FITC, Cy3, Cy5, Cy7). After imaging of the last sequencing cycle, the probes were stripped off one more time and samples were imaged to receive the autofluorescent background of the tissue in each channel for background correction.

### Imaging

Imaging was performed using a digital slide scanner (Slideview VS200, Olympus, Tokio, Japan) connected to external LED source (Excelitas Technologies, X-Cite Xylis, Mississauga, Canada). Fluorescence filter cubes and wheels were equipped with a pentafilter (AHF, excitations: 352-404 nm, 460-488 nm, 542-566 nm, 626-644 nm, 721-749 nm; emissions 416-452 nm, 500-530 nm, 579-611 nm, 665-705 nm, 767-849 nm). The images were obtained with a sCMOS camera (2304 × 2304, ORCA-Fusion C14440-20UP, 16 bit, Hamamatsu, Japan), and Olympus universal-plan super apochromat 40× (0.95 NA/air, Olympus). For each slide and cycle an image in DAPI, FITC, Cy3, Cy5 and Cy7 was taken, and the scanned areas were outlined and saved in order to perform repetitive cycle imaging. Extended focus imaging (EFI) was used to automatically discard unfocused z-stack images, resulting in bright and focused *in situ* signals.

### Image analysis

Imaging data was analysed with a custom pipeline provided by Cartana (part of 10x Genomics) and published pipelines found in the repository (https://github.com/Moldia/HybrISS) that handles image processing and gene calling [18]. All code was written in MATLAB and additionally, a CellProfiler (v.2.1.1) pipeline with the MultiStackReg, StackReg and TurboReg plugins was used [19]. Therefore, images from all sequencing cycles were exported into.tiff-format and arranged through alignment of the DAPI channel of the first sequencing cycle with the channels of each sequencing cycle. Then, images were split into multiple smaller images to allow analysis in CellProfiler.

As each fluorescent colour has different intensity values for RCP signals in their respective colours, we normalized the intensity values to 10 000 and calculated thereby a multiplication factor. E.g.: the median intensity of RCP signal in Cy5 was 5 000 and need to be multiplied by two, to reach 10 000. This multiplication factor was calculated for each fluorescent colour and then used to normalise the median intensity of all RCP signals. The received multiplication factor for each channel was integrated in the CellProfiler pipeline and additionally, the background of each channel was subtracted from each sequencing cycle to reduce the autofluorescence of the tissue. A pseudo-general stain was created by combining the 4 readout detection probe channels of the first sequencing cycle into one merged image. Additionally, a pseudo-anchor for each sequencing cycle was generated to perform a second, more exact alignment to the pseudo-general stain. The RCPs of the pseudo-general stain were detected to obtain the x- and y–coordinates of the ISS genes. Based on these positions, the fluorescence intensities in each of the 4 channels (FITC, Cy3, Cy5 and Cy7) were measured. This procedure was performed for all sequencing cycles to derive the measured intensities. The highest intensity value in each sequencing cycle was then assigned as real signal and further used for decoding with MATLAB [18]. For the verification of the signals, the selected transcripts were plotted on a DAPI-stained image [18].

### Compartment building by morphology and virtual H&E

To combine the advantage of ISS and H&E morphology on the exact same tissue section, virtual H&E images were created of the ISS hybridised tissue sections based on DAPI and FITC images, as described by Giacomelli et al. [20]. Virtually stained H&E images of the patient tissue samples were evaluated by a pathologist specialised on colon cancer, who then assigned tissue areas into neoplastic and non-neoplastic areas (“morphology-based” approach). Afterwards, the outlines of the neoplastic and non-neoplastic areas were digitalised and converted into binary tissue compartments as described in greater detail in the supplementary material.

### Tissue compartment building by gene expression

Most of the via ISS analysis detected transcripts were expressed in neoplastic-and non-neoplastic tissue compartment. However, specific genes showed clear overexpression of their transcripts in neoplastic vs. non-neoplastic tissue compartments in all analysed tissue samples. Based on this observation, we evaluated if such a set of genes can be used to automatically classify tissue into neoplastic and non-neoplastic tissue compartments without future need of histological information. In doing so, *in situ* signals of the respective genes were computationally represented as dots of a certain size, and computationally superimposed to form connected areas, as shown in Fig. 1. The detailed description of this approach can be found in the supplement materials. In short, the dot like signals were expanded by 50-180 pixels and thereby merging and forming larger and connected areas. In order to remove small gaps in connected tissue compartments, the python library openCV [21] was used. A threshold technique was then applied to convert this superimposition into a binary neoplastic tissue compartment. In the next step, the overlap between the “morphological-based” and “gene expression-based” binary neoplastic tissue compartment was calculated for each sample. Further, the sample overlaps were combined via geometric mean to a mean overlap value that was used to rate the set. The mean overlap values were calculated for alternating compositions of genes. In the end, the set of ISS genes that achieved the highest mean overlap value was selected and used for the statistical testing. The binary non-neoplastic tissue compartment of a patient sample was obtained by excluding the neoplastic one from the entire tissue compartment, where latter was derived by superimposition of the dot representations of all detected ISS genes and all cell nuclei.

**Figure 1:**
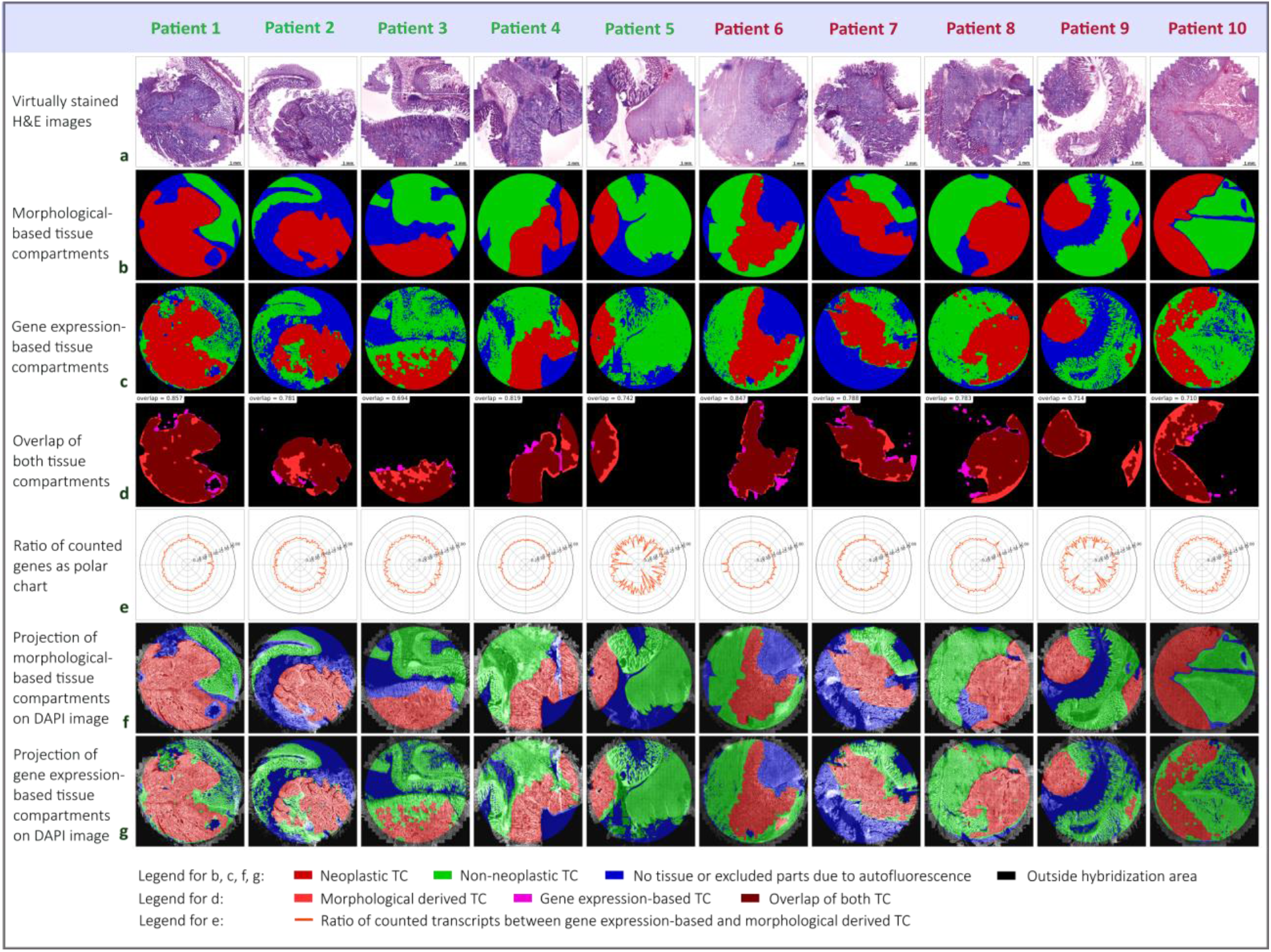
Generation of expression-based tissue compartments and overlap with morphological tissue compartments. **a)** The virtually stained H&E images of the samples from non-relapsed (patient 1-5) and relapsed patients (patient 6-10). **b)** Tissue classified into neoplastic and non-neoplastic tissue compartment by a pathology expert based on morphological characteristics. **c)** Gene expression-based neoplastic and non-neoplastic tissue compartment by using the *in situ* sequencing tumour gene signature (*EREG, MET, BIK, CD44, ITGAV, MYBL2, CCND1* and *S100A4*). **d)** Overlap of the morphological- and the gene expression-based tissue compartment for neoplastic tissue. The mean overlap value for the tumour gene signature is 0.77. **e)** Ratios of the counted gene per cell value between the gene expression-based and the morphological- based neoplastic tissue compartment depicted as polar chart. Thereby, each data point shows the ratio for a certain *in situ* sequencing gene. **f)** Projection of morphological obtained tissue compartment on the DAPI images. **g)** Projection of gene expression-based tissue compartment on the DAPI images. Size bar is the same for all images.

### Gene counting and statistical testing

In order to identify significances in the distribution of genes of a certain type in the binary neoplastic and non-neoplastic tissue compartment, the number of genes are counted within both tissue compartments. We developed a script to create the tissue compartments and quantify the gene counts, namely genes-to-count (GTC-Tool) available at the repository (https://github.com/spatialhisto/GTC). To take into account differences in sizes and cell numbers of the tissue compartments, counts were normalised by the number of detected cell nuclei within a tissue compartment (see supplementary materials). A two-tailed paired t-test is used for the statistical testing of differences of the gene per cell values in the neoplastic and the non-neoplastic tissue compartment in the patient distribution. Further, a two-tailed independent t-test is applied for statistical testing of the gene per cell value in the neoplastic tissue compartment in both patient distributions, the relapsed and the non-relapsed group. The statistical testing is done with a significance level α/2=0.05 for the morphological-based as well as for the gene expression-based tissue compartment.

## Results

### Colon tissue can automatically be classified by dRNA-HybISS based gene expression data into neoplastic and non-neoplastic compartments

The *in situ* sequencing analysis was performed on tissue samples containing neoplastic and non-neoplastic tissue including stroma and/or healthy colonic epithelium. To identify neoplastic or non-neoplastic compartments, we developed a gene expression-based tool called genes-to-count (GTC-tool), based on histopathological expertise. Thereby a set of signature genes which are highly expressed in neoplastic tissue served as template for defining a neoplastic tumour compartment. To do so, virtually stained H&E images produced of each tumour (Fig. 1a) were annotated by a board certified pathologist specialised on colon cancer to classify neoplastic and non-neoplastic tissue compartments based on the morphology of the tissues (Fig. 1b). The gene expression-based neoplastic and non-neoplastic tissue compartments were then generated based on specific expression patterns of a 8-gene set containing the genes *EREG, MET, BIK, CD44, ITGAV, MYBL2, CCND1* and *S100A4*, we refer to as tumour gene signature. This tumour gene signature achieved the highest mean overlap of 77% for neoplastic tissue compartment between morphological- and expression-based tissue compartments (Fig 1c). The remaining tissue was defined as non-neoplastic tissue. Thereby the GTC-tool integrates the spatial coordinates of each decoded transcript and nucleus into its tissue compartments and automatically quantify RNA transcripts in the respective compartment. However, small areas of some patient samples, i.e. of patients 2, 6, 7 and 9, cannot be used for analysis due to tissue loss/damage during the sequencing procedure or high autofluorescence and were thus excluded, as shown in Fig. 1a.

The overlap of the morphological-based and gene expression-based neoplastic tissue compartment is shown in Fig. 1d for each patient sample (patient 1=85.7%, patient 2=78.1%, patient 3=69.4%, patient 4=81.9%, patient 5=74.2%, patient 6=84.7%, patient 7=78.8%, patient 8=78.3%, patient 9=71.4% and patient 10=71.0%).

The ratio of the morphological-based and gene expression-based neoplastic tissue compartment gene per cell counts are shown as polar chart in Fig. 1e. As can be seen therein, only a slight variation occurs between both neoplastic tissue compartments. For a better visualization regarding the localization of the compartments within the tissue architecture, DAPI-images were superimposed with the morphological-based (Fig 1f) and the expression-based tissue compartment (Fig 1g).

### Spatially differential gene expression in neoplastic vs. non-neoplastic tissues

The expression level of each transcript was examined by comparing its counts per cell in the neoplastic vs. the non-neoplastic tissue compartments in 10 colon cancer patients (Fig. 2).

**Figure 2:**
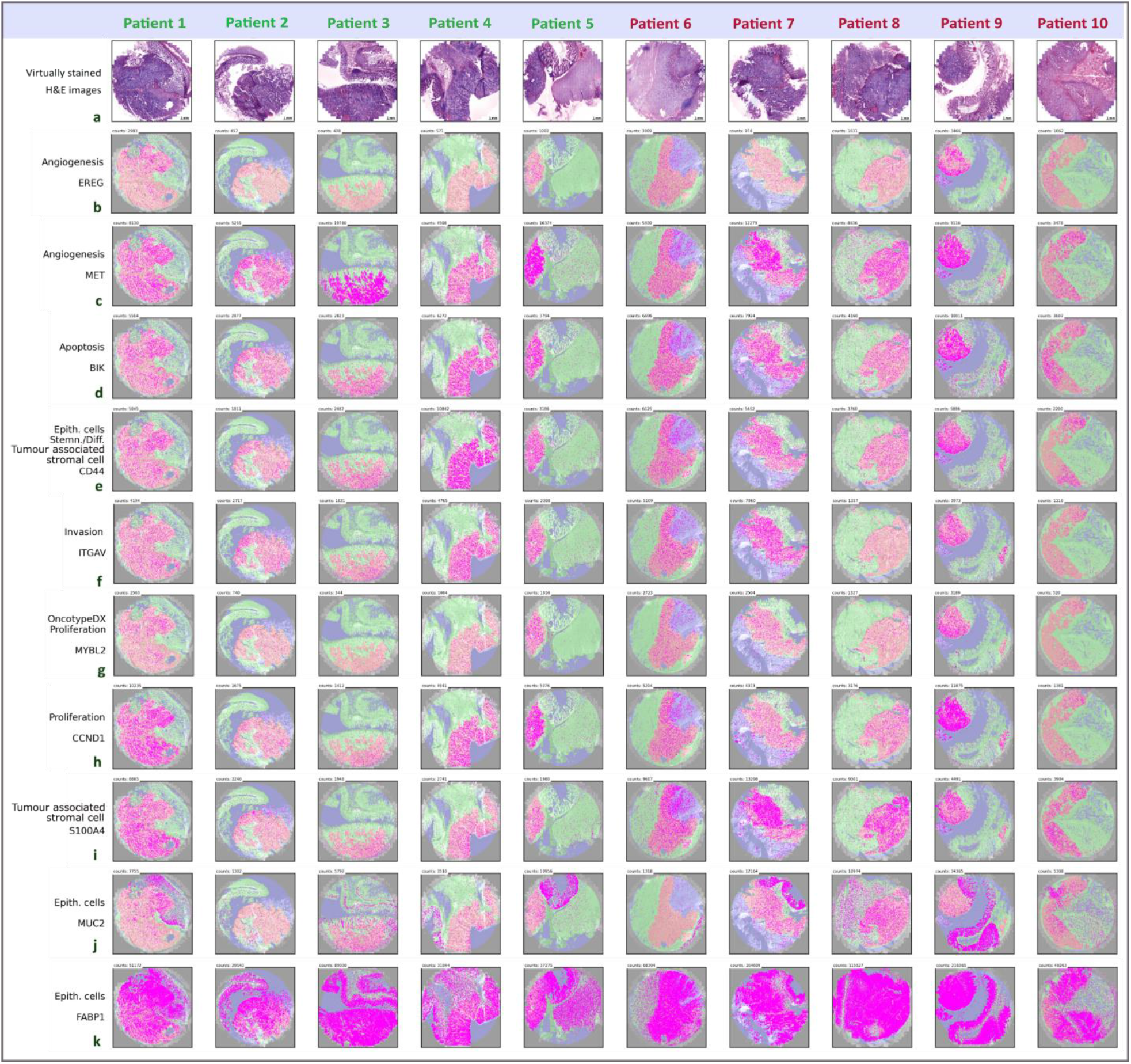
Spatial distribution of 10 selected genes in neoplastic and non-neoplastic tissue. **a)** The virtually stained H&E images of the samples from non-relapsed (patient 1-5) and relapsed patients (patient 6-10). **b-i)** The tumour gene signature was used for the creation of the neoplastic tissue compartment. **j)** Exemplified expression and the spatial distribution of *MUC2, a gene expressed in non-neoplastic epithelial- and cancer cells*. **k)** Exemplified expression and the spatial distribution *FABP1*, a high expressed gene in colonic tissue. Total counts of each transcript are depicted in each image and size bar is the same for all images.

Therefore, volcano plots with a significance level α/2=0.05 were generated to define significantly upregulated genes associated with different biological processes (Fig. 3). 98 significantly upregulated genes were identified in the expression-based tissue compartment (Fig 3c, d), whereas 81 genes were significantly upregulated in the morphological-based tissue compartment (Fig. 3d, e). All significantly upregulated genes detected in the morphological tissue compartment (81 of 81), could also be detected in the expression-based tissue compartment, additionally 17 genes showed an elevated expression in the expression-based tissue compartment, which can be seen highlighted in red in Fig. 3c. In the morphological neoplastic tissue compartment (Fig. 3a), highly significant upregulated genes and/or genes with high fold changes were observed: apoptosis related genes (*CASP3, BIK*), genes associated with proliferation (*CCND1, PCNA, MYBL2*), tumour associated stromal genes (*TIMP1, CXCL1, COL1A2, S100A4, CD44*),marker genes for energy metabolism (*LDHA, PKM*) and oxidative stress (*SOD1, GPX1, PRDX2*), stemness-(*CD44*) and angiogenesis-(*MET*) associated genes, and Oncotype DX genes (*INHBA, MYBL2*).

**Figure 3:**
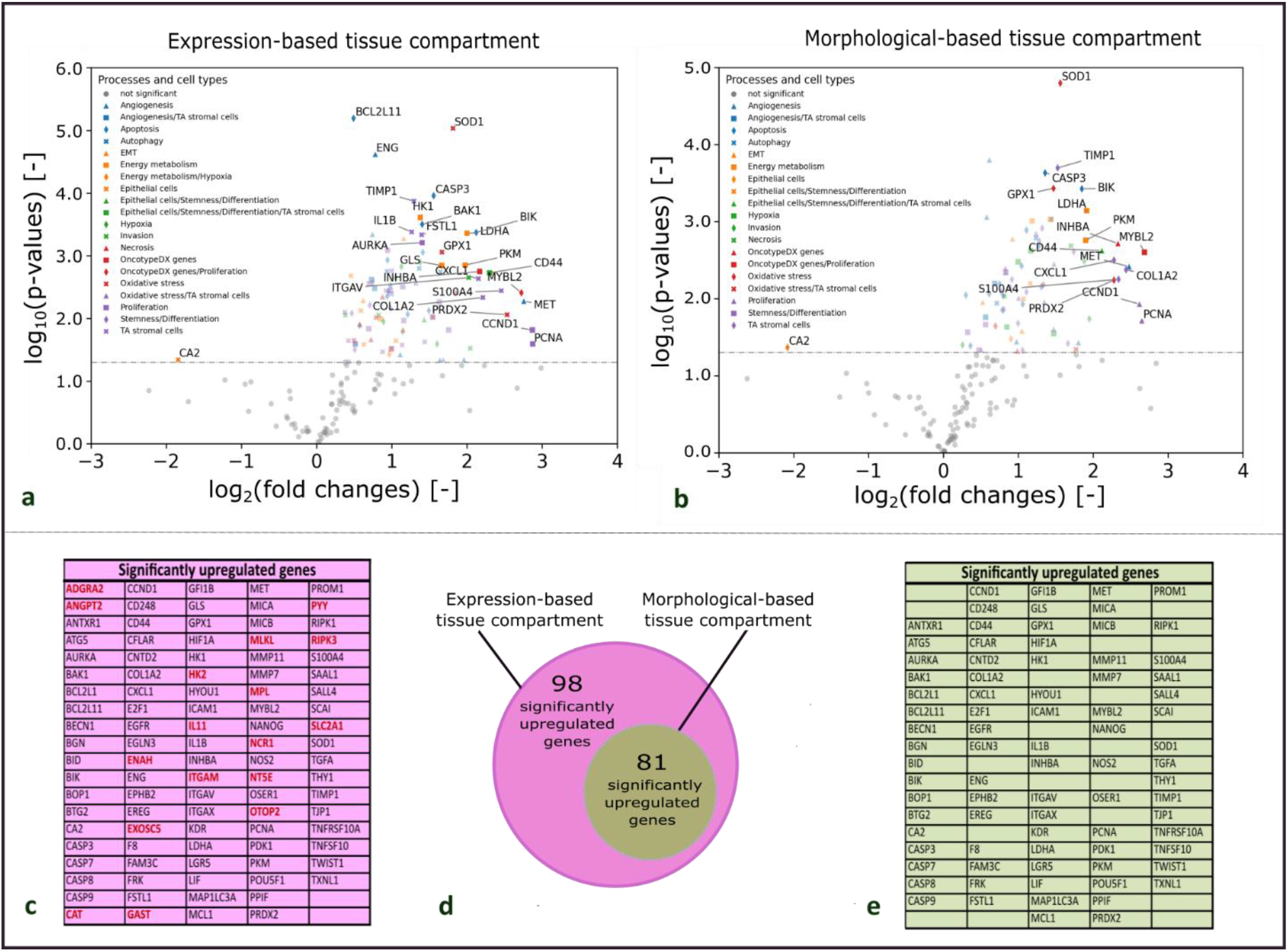
Significantly upregulated genes in neoplastic vs. non-neoplastic tissues compartments (N=10). **a, b)** Volcano plot of upregulated genes in the expression-based tissue compartment, and morphological-based tissue compartment. Genes which show a high significance and/or high fold change between the neoplastic and non-neoplastic tissue compartments are labelled by name. Genes belonging to different biological processes are marked with different symbols in different colours to achieve an overview of relevant processes upregulated in neoplastic tissue compartments. Each dot represents an individual gene, a two-sided paired t-test is used for statistical testing with a significance level α/2=0.05 (horizontal line). **c)** List of all significantly upregulated genes in the expression-based tissue compartment. Red labelled genes were only found significantly differential expressed in the expression-based tissue compartment. Black labelled genes are concordant between expression- and morphological-based tissue compartments. **d)** Diagram of the amount of genes upregulated in the expression-based and the morphological tissue compartment. **e)** List of all significantly upregulated genes in the morphological tissue compartment. TA stromal cells = tumour associated stromal cells, EMT = epithelial-mesenchymal transition.

In the expression-based neoplastic tissue compartment (Fig. 3b) the following genes indicated a high-significantly increase in expression and/or a high fold change: apoptosis related genes (*BCL2L11, ENG, CASP3, BAK, BIK*), genes associated with proliferation (*AURKA, CCND1, PCNA, MYBL2*) and tumour associated stromal genes (*IL1B, FSTL1, TIMP1, CXCL1, COL1A2, S100A4, CD44*), marker genes for energy metabolism (*HK1, GLS, LDHA, PKM*) and oxidative stress (*SOD1, GPX1, PRDX2*), stemness-(*CD44*),invasion-(*ITGAV*) and angiogenesis-(*ENG, MET*) associated genes as well as Oncotype DX genes (*INHBA, MYBL2*). All significantly expressed genes in the expression-based compartment are listed in Fig. 3c, whereas the list of genes upregulated in the morphological-based compartment can be seen in Fig. 3e.

### Upregulated genes in the neoplastic tissue compartment of relapsed patients vs. non-relapsed patients

To investigate the differences of gene expression in the neoplastic tissue compartment of 5 relapsed patients in comparison to 5 non-relapsed stage II colon cancer patients, a two-sided independent t-test with a significance level α/2=0.05 was performed. The outcome is illustrated in Fig. 4a by a volcano plot. Three genes were identified showing a significant increase of expression in relapsed patients compared to non-relapsed. The expression level of *OTOP2* (Fig. 4b) indicate a significant upregulation with 2.6 counts per 1000 cells in relapsed compared to 1.9 counts per 1000 cells for non-relapsed patients (p-value=0.0042). The expression for the transcript *FGFR2* (Fig. 4c) showed 4.9 counts per 1000 cells for relapsed and 3.1 for non-relapsed patients (p-value=0.0137). For *MMP11* significantly elevated expression levels for relapsed patients were observed with 49.2 counts per 1000 cells for relapsed in comparison to 16.4 counts per 1000 cells for non-relapsed patients (p value=0.0415).

**Figure 4:**
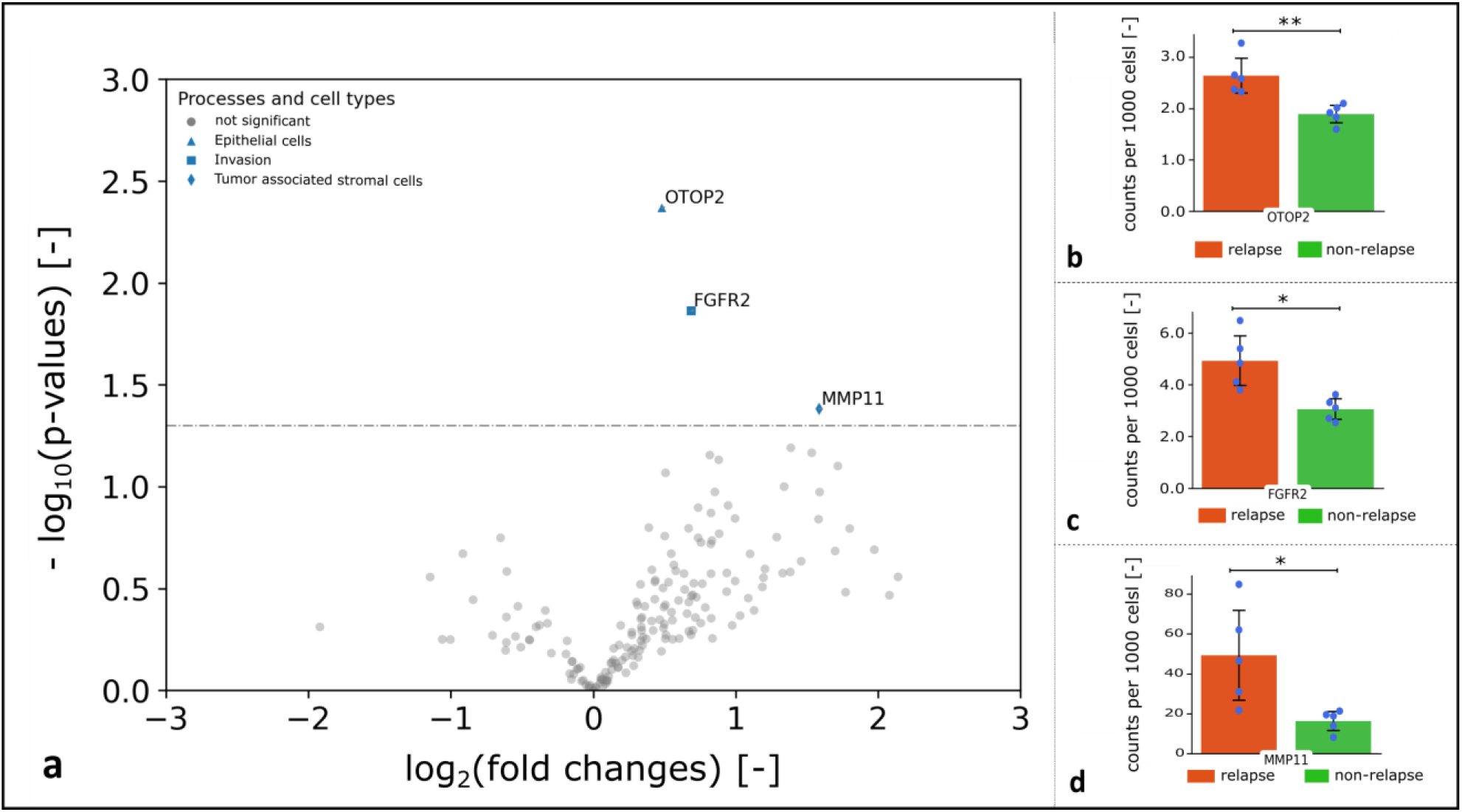
Upregulated genes in neoplastic tissue compartments in relapsed patients in comparison to non-relapsed patients (N=10). **a)** Volcano plot with a significance level α/2=0.05 of significantly upregulated genes in the neoplastic tissues compartment of relapsed patients in comparison to non-relapsed patients. **b-d)** The expression level of *OTOP2, FGFR* and *MMP11* in relapsed patients (orange) indicated a significant increase in comparison to non-relapsed patients (green). Significant differences (*p < 0.05 and **p < 0.005) were highlighted with bars and asterisks.

## Discussion

In the current study, we applied for the first time a direct RNA ISS approach in colon cancer with sub-cellular resolution investigating the spatial expression of 175 genes. In stage II colon cancer, we identified *FGFR2, MMP11* and *OTOP2* as upregulated genes in the tumour compartment in patients which relapsed. Importantly, *FGFR2* and *MMP11* are druggable targets in other cancer entities and could become novel predictive biomarkers in stage II colon cancer. Moreover, we developed a genes-to-count (GTC) tool to accurately classify colon tissue into neoplastic and non-neoplastic compartments using an 8-gene tumour gene signature and quantify spatial gene expression. The spatial ISS approach allows to yield novel insights into predictive CRC biomarkers beyond bulk tissue sequencing.

We observed a significantly elevated expression of the Fibroblast Growth Factor Receptor 2 (*FGFR2*) in tumour compartments of relapsed colon cancer stage II patients. This transcript has been identified as a potential therapeutic target for CRC by Matsuda et al. and Li et al. as upon binding to ligands a series of downstream signalling pathways are activated that are involved in differentiation, survival and proliferation and play major roles in the progression of CRC [22, 23]. Interestingly *FGFR2* has been shown to be druggable in other tumour entities. Pemigatinib and erdafitinib, two anti-*FGFR* agents, are already approved by the Food and Drug Administration (FDA) for treatment of cholangiocarcinoma and urothelial carcinoma, and various *FGFR* inhibitors are currently being evaluated in preclinical and clinical trials [24]. Due to overexpression of *FGFR2* in numerous tumours and its significant role in progression and tumourigenesis, *FGFR2* is a promising target for treatment of stage II colon cancer patients in future [25]. Matrix metalloproteinase 11 (*MMP11*) belongs to the family of zinc dependent endopeptidases and unlike other MMPs, *MMP11* is having some unique characteristics [26]. *MMP11* is secreted in an enzymatically active form, while most other MMPs are released as inactive enzymes. Additionally it is promoting the signal transduction of protein kinase B (*AKT*) / Forkhead box protein O1 (*FoxO1*) / insulin-like growth Factor-1 (I*GF1*), which is associated with the lysis of collagen type VI and proliferation of connective tissue around the stroma in cancerous tissues [26, 27]. These features indicate that *MMP11* plays a unique role in the development of malignant tumours, their progression and metastasis [26]. A study performed by Zhuang et al. showed that *MMP11* was associated with various signalling pathways involved in tumour development in breast cancer and high expression of *MMP11* is associated with poor prognosis for patients [28]. Yang et al. identified *MMP11* as a key cancer driver in lung adenocarcinoma and a potential target for antibody therapy as application of anti-*MMP11* antibodies suppressed the growth of tumours in xenograft models [29]. In contrast to *FGFR2* and *MMP11*, less published data is available for Otopetrin 2 (*OTOP2*) in CRC. *OTOP2* encodes a proton-selective channel, transferring protons into the cell cytosol in response to low pH in various epithelia [30]. Recently, scRNAseq analysis identified a new subtype of cells within the intestinal crypts, being positive for *OTOP2* and *BEST4* (calcium-sensitive chloride channel), namely *BEST4/OTOP2* cells, eventually responsible for electrolyte transportation [31]. In colorectal cancer and inflammation, loss of *BEST4/OTOP2* cells have been described [31]. In our study we overserved that *OTOP2* is overexpressed in relapsed stage II colon cancer patients. In contrast, Qu et al and Guo et al showed that elevated levels of *OTOP2* in cell line experiments is lowering tumour proliferation and high expression of *OTOP2* in bulk CRC tissues was significantly correlated with better overall survival. It is importantly to mention, that the CRC cohort of Guo et al did not focus on stage II colon cancer and relied on bulk RNA expression data [32]. Therefore, comparison between our cohort of stage II colon cancer and a broader cohort of CRC tissue and different stages is hampered. Moreover, bulk RNA analysis generally includes tumour and stroma tissue resulting in an average gene expression, whereas spatial expression analysis enables us to only address the neoplastic tissue compartment allowing a less diluted observation [33].

Evaluation of tumour tissues and their histological compartments, such as tumour, stroma and immune cells need histological know-how and experience. Several powerful AI based tools are evolving, but usually need large training sets to identify the respective tissue compartments [3]. Instead of AI based tissue classification, we make use of spatial ISS data to define tumour compartments, i.e. genes which are highly expressed in neoplastic colon tissues allows us to generate tissue compartments of neoplastic tissue. A major advantage of the resulting compartments is that they are independent from tissue histology as they rely only on gene expression and can therefore be applied to other colon cancer samples without the need of large training image data sets. Similar approach was developed by Meylan et al, who identified tertiary lymphoid structures in renal cell carcinomas based on a 29-gene signature dominated by immune globulin genes, but with lower resolution of a bin size of 55μm using Visium spatial transcriptomics (10x Genomics) [34]. When comparing our data of both approaches, I.e. morphology-based vs. expression-based, we observed that the expression-based approach shows more granular and precise representation of neoplastic tissue in three patient samples (patient 2, 3 and 10), especially at tumour border regions. Both approaches, morphological- and expression-based tissue compartments showed high concordance, also in the context of differential gene expression of neoplastic vs. non-neoplastic tissue. All 81 upregulated genes by using the morphological-based approach (neoplastic vs. non-neoplastic tissue compartments) were also identified in the expression-based approach, confirming the equally good performance and precision of the created expression-based tissue compartments. In addition, 17 more differentially expressed genes were found (total of 98) by the use of the expression-based approach.

Recognizing the extraordinary potential of spatial transcriptomic datasets in revealing detailed cellular- and tissue organization, data analysis remains challenging. A multitude of analysis and visualization tools for pre-processing, clustering, cell phenotyping, and cell-cell interaction are being developed continuously but no gold standard has crystallised yet [35]. For example, segmentation of cells and assigning expressed transcripts to identify the underlying cell type can be performed by sophisticated methods such as Baysor [36], JSTA [37] or modified pipelines from pciSeq [38] and Scanpy [39, 40] among many others. In our data the quality of ISS results strongly depend on tissue characteristics such as autofluorescence or good fixation of tissue, as optimal sensitivity and specificity of the ISS methodology requires bright, clear signals and low background [41]. Highly autofluorescent tissue structures such as elastin and collagen [42] can occlude or mimic *in situ* signals and would result in wrong base calling during decoding. Therefore, we have developed and included a background subtraction step, to reduce high autofluorescence especially for channels detected in shorter wavelengths such as FITC and Cy3. We observed an estimated ~25% higher number of correctly assigned transcripts vs. without background subtraction. Another obstacle in spatial transcriptomic data sets is gene expression normalization between samples. Available tools such as scran [43] or SCnorm [44] are inspired by scRNAseq studies but there is no “one-size-fits-all” solution [39, 33] and were not developed for sub-cellular ISS data sets. As dRNA-HybISS yield subcellular resolution, we could normalise our expression data for each individual tumour sample. To do so, we segmented cells using available scripts (provided by Cartana) and normalised the number of transcripts to cell counts. For our aim to investigate the spatial tissue composition of colon cancer stage II this approach is feasible and similar to previously described approaches [46].

## Conclusion

In conclusion, *in situ* sequencing revealed novel potential predictive biomarkers in colon cancer stage II, namely *MMP11, FGFR2* and *OTOP2*, relevant for relapse of disease. Our developed and open-access available GTC-tool allows to accurately capture the tumour compartment and quantify gene expression in colon cancer tissue.

## Acknowledgements

KS was supported by the Doctoral School “Translational Molecular and Cellular Biosciences” of the Medical University of Graz and LB was supported by the PhD Program AMBRA (Advanced medical biomarker research) of the Medical University of Graz together with the FFG K1 center CBmed (Center for Biomarker Research in Medicine). This work was supported by the K1 COMET Competence Center CBmed, which is funded by the Federal Ministry of Transport, Innovation and Technology (BMVIT); the Federal Ministry of Science, Research and Economy (BMWFW), Land Steiermark (Department 12, Business and Innovation), the Styrian Business Promotion Agency (SFG), and the Vienna Business Agency. The COMET program is executed by the Austrian Research Promotion Agency (FFG). The authors thank Daniel Kummer and Martina Tomberger for technical support. The tissue samples used in this project have been provided by Biobank Graz.

## Abbreviations

CRC: colorectal cancer
Cy3: Cyanine 3
Cy5: Cyanine 5
Cy7: Cyanine 7
DAPI: 4,6-diamidino-2-phenylindole
dRNA-HybISS: direct RNA - Hybridisation *based in situ* sequencing
EMT: epithelial-mesenchymal transition
FFPE: formalin-fixed, paraffin-embedded
FITC: Fluorescein isothiocyanate
H&E: Hematoxylin and eosin stain
ISS: *in situ* sequencing
HybISS: Hybridisation *based in situ*sequencing LED: light-emitting diode
RCA: rolling circle amplification
RCP: rolling circle product
RNA: Ribonucleic acid
scRNA: single cell RNA sequencing
TC: tissue compartment

