## Supplemental material for "Spatial tumour gene signature discriminates neoplastic from non-neoplastic compartments in colon cancer: unravelling predictive biomarkers for relapse"

### 1. List of gene panel

*Table 1:* List of designed gene panel

| Biological process/group | Subgroups | Gene (according to Human protein atlas [1,2]) | ENS number (derived from Ensembl genome browser) |
|---|---|---|---|
| **Angiogenesis** | | ANGPT1 | ENSG00000154188 |
| | | ANGPT2 | ENSG00000091879 |
| | | FLT4 | ENSG00000037280 |
| | | KDR (VEGFR-2) | ENSG00000128052 |
| | | TIE1 | ENSG00000066056 |
| | | EREG | ENSG00000124882 |
| | | ENG (CD105) | ENSG00000106991 |
| | | MET | ENSG00000105976 |
| | | CD248 (TEM1) | ENSG00000174807 |
| | | ADGRA2 (TEM5) | ENSG00000020181 |
| | | PLXDC1 (TEM7) | ENSG00000161381 |
| | | ANTXR1 (TEM8) | ENSG00000169604 |
| | | TEK | ENSG00000120156 |
| | | F8 | ENSG00000185010 |
| **Apoptosis** | **Pro-apoptotic** | BBC3 (PUMA) | ENSG00000105327 |
| | | TNFSF10 (TRAIL) | ENSG00000121858 |
| | | BID | ENSG00000015475 |
| | | BIK | ENSG00000100290 |
| | | BCL2L11 (Bim) | ENSG00000153094 |
| | | BAK1 | ENSG00000030110 |
| | **Inhibitors** | CFLAR (cFLIP) | ENSG00000003402 |

| | | | |
|---|---|---|---|
| | | BCL2L1 (BCLX) | ENSG00000171552 |
| | | MCL1 | ENSG00000143384 |
| | **Receptors** | TNFRSF10A (DR4) | ENSG00000104689 |
| | | TNFRSF10B (DR5) | ENSG00000120889 |
| | **Caspases** | CASP8 | ENSG00000064012 |
| | | CASP9 | ENSG00000132906 |
| | | CASP7 | ENSG00000165806 |
| | | CASP3 | ENSG00000164305 |
| **Autophagy** | | MAP1LC3A (LC3) | ENSG00000101460 |
| | | ATG5 | ENSG00000057663 |
| | | BECN1 (Beclin 1) | ENSG00000126581 |
| **Necrosis** | | RIPK3 | ENSG00000129465 |
| | | MLKL | ENSG00000168404 |
| | | HMGB1 | ENSG00000189403 |
| | | RIPK1 | ENSG00000137275 |
| | | PPIF | ENSG00000108179 |
| **Proliferation** | **Proliferation** | PCNA | ENSG00000132646 |
| | | MCM2 | ENSG00000073111 |
| | | CNTD2 | ENSG00000105219 |
| | | EXOSC5 | ENSG00000077348 |
| | | E2F1 | ENSG00000101412 |
| | | MYBL2 | ENSG00000101057 |
| | | CCND1 | ENSG00000110092 |
| | | CCNE1 | ENSG00000105173 |
| | | GRB7 | ENSG00000141738 |
| | | RPS6KB1 | ENSG00000108443 |
| | | AURKA | ENSG00000087586 |
| | | SAAL1 | ENSG00000166788 |
| | | TGFA | ENSG00000163235 |
| | | EGFR | ENSG00000146648 |
| | | KIT | ENSG00000157404 |
| | | URGCP | ENSG00000106608 |

| | | | |
|---|---|---|---|
| | **Inhibitors** | BOP1 | ENSG00000261236 |
| | | BTG2 | ENSG00000159388 |
| | | FRK | ENSG00000111816 |
| **Oxidative stress** | | SOD1 | ENSG00000142168 |
| | | GPX1 | ENSG00000233276 |
| | | CAT | ENSG00000121691 |
| | | GSR | ENSG00000104687 |
| | | NOS2 | ENSG00000007171 |
| | | TXNL1 | ENSG00000091164 |
| | | PRDX2 | ENSG00000167815 |
| | | OSER1 | ENSG00000132823 |
| **Hypoxia** | | HIF1A | ENSG00000100644 |
| | | HIF3A | ENSG00000124440 |
| | | EGLN3 | ENSG00000129521 |
| | | PDK1 | ENSG00000152256 |
| | | SLC2A1 (GLUT1) | ENSG00000117394 |
| | | HYOU1 | ENSG00000149428 |
| **Stemness/Differentiation** | | ALDH1A1 | ENSG00000165092 |
| | | NANOG | ENSG00000111704 |
| | | SALL4 | ENSG00000101115 |
| | | CD44 | ENSG00000026508 |
| | | PROM1 (CD133) | ENSG00000007062 |
| | | BMI1 | ENSG00000168283 |
| | | MPL (CD110) | ENSG00000117400 |
| | | EPHB2 | ENSG00000133216 |
| | | LGR5 | ENSG00000139292 |
| | | IL6 | ENSG00000136244 |
| | | POU5F1 | ENSG00000204531 |
| | | SOX2 | ENSG00000181449 |
| **Invasion** | | LAMC2 (Laminin-5, γ2) | ENSG00000058085 |
| | | ITGAV | ENSG00000138448 |
| | | L1CAM | ENSG00000198910 |

| | | | |
|---|---|---|---|
| | | MMP7 | ENSG00000137673 |
| | | ENAH (MENA) | ENSG00000154380 |
| | | FGFR2 | ENSG00000066468 |
| | | SCAI | ENSG00000173611 |
| | | MIEN1 | ENSG00000141741 |
| | | TIAM1 | ENSG00000156299 |
| **Epithelial-Mesenchymal Transition** | | TJP1 | ENSG00000104067 |
| | | FN1 (Fibronectin ) | ENSG00000115414 |
| | | TWIST1 | ENSG00000122691 |
| | | FOXC2 | ENSG00000176692 |
| | | ZEB1 | ENSG00000148516 |
| | | FAM3C | ENSG00000196937 |
| **Energy metabolism** | | SLC2A1 (GLUT1) (already in table 1, other biological process) | ENSG00000117394 |
| | | HK1 | ENSG00000156515 |
| | | HK2 | ENSG00000159399 |
| | | PKM2 | ENSG00000067225 |
| | | LDHA | ENSG00000134333 |
| | | GLS | ENSG00000115419 |
| | | GLUD1 | ENSG00000148672 |
| **OncotypeDX genes** | **Cell cycle genes** | MYBL2 (already in table 1, other biological process) | ENSG00000101057 |
| | **Stromal genes** | BGN | ENSG00000182492 |
| | | INHBA | ENSG00000122641 |
| | **Early response gene** | GADD45B | ENSG00000099860 |
| **Epithelial cells** | **Colonocytes (Enterocytes)** | ANPEP | ENSG00000166825 |
| | | CA1 | ENSG00000133742 |

| | | | |
|---|---|---|---|
| | | CA2 | ENSG00000104267 |
| | | FABP1 | ENSG00000163586 |
| | | BEST4 | ENSG00000142959 |
| | | OTOP2 | ENSG00000183034 |
| | | GUCA2B | ENSG00000044012 |
| | | SLC26A3 | ENSG00000091138 |
| | **Goblet cells** | MUC2 | ENSG00000198788 |
| | | MUC5AC | ENSG00000215182 |
| | | SPDEF | ENSG00000124664 |
| | | KLF4 | ENSG00000136826 |
| | | TFF1 | ENSG00000160182 |
| | | TFF3 | ENSG00000160180 |
| | **Enteroendocrine cells** | CHGA | ENSG00000100604 |
| | | TBXT | ENSG00000164458 |
| | | ZGLP1 | ENSG00000220201 |
| | | GIP | ENSG00000159224 |
| | | SST | ENSG00000157005 |
| | | NTS | ENSG00000133636 |
| | | PYY | ENSG00000131096 |
| | | GAST | ENSG00000184502 |
| | | MLN | ENSG00000096395 |
| | **Tuft cells** | DCLK1 | ENSG00000133083 |
| | | TRPM5 | ENSG00000070985 |
| | | POU2F3 | ENSG00000137709 |
| | | GFI1B | ENSG00000165702 |
| | **Stem cells** | OLFM4 | ENSG00000102837 |
| | | CD44 (already in table 1, other biological process) | ENSG00000026508 |
| | | PROM1 (CD133) (already in table 1, other biological process) | ENSG00000007062 |

| | | | | |
|---|---|---|---|---|
| | | | LGR5 (already in table 1, other biological process) | ENSG00000139292 |
| | | | ALDH1A1 (already in table 1, other biological process) | ENSG00000165092 |
| **Tumor-associated stromal cells (TASC)** | **Cancer- associated fibroblasts (CAFs)** | **Mesenchymal stem cell-like** | CD105 (already in table 1, other biological process) | ENSG00000106991 |
| | | | THY1 (CD90) | ENSG00000154096 |
| | | | NT5E (CD73) | ENSG00000135318 |
| | | | CD44 (already in table 1, other biological process) | ENSG00000026508 |
| | | **Endothelial-like** | ICAM1 (CD54) | ENSG00000090339 |
| | | | TEK (TIE2) | ENSG00000120156 |
| | | | VCAM1 (CD106) | ENSG00000162692 |
| | | | CDH5 (CD144) | ENSG00000179776 |
| | | **MyoFibroblast-like** | TNC | ENSG00000041982 |
| | | | TAGLN | ENSG00000149591 |
| | | | PDGFA | ENSG00000197461 |
| | | | TGFB3 | ENSG00000119699 |
| | | **Pericyte-like** | CSPG4 (NG2) | ENSG00000173546 |
| | | **Matrix remodelling** | S100A4 (FSP1) | ENSG00000196154 |
| | | | MMP2 | ENSG00000087245 |

| | | | | |
|---|---|---|---|---|
| | | | DCN | ENSG00000011465 |
| | | | COL1A2 | ENSG00000164692 |
| | | **others** | FSTL1 | ENSG00000163430 |
| | | | TIMP1 | ENSG00000102265 |
| | | | LIF | ENSG00000128342 |
| | | | IL11 | ENSG00000095752 |
| | **Cancer-associated adipocytes (CAAs)** | | IGFBP2 | ENSG00000115457 |
| | | | MMP11 | ENSG00000099953 |
| | | | IL6 (already in table 1, other biological process) | ENSG00000136244 |
| | | | IL1B | ENSG00000125538 |
| | | | IL8 | ENSG00000169429 |
| | | | FABP4 | ENSG00000170323 |
| | **Cancer-associated endothelial cells (CAECs)** | | S100A4 (FSP1) (already in table 1, other biological process) | ENSG00000196154 |
| | | | CXCL1 | ENSG00000163739 |
| | | | CD248 (TEM1) (alredy in list) | ENSG00000174807 |
| | | | ADGRA2 (TEM5) (already in table 1, other biological process) | ENSG00000020181 |
| | | | PLXDC1 (TEM7) (already in table 1, other biological process) | ENSG00000161381 |
| | | | ANTXR1 (TEM8) | ENSG00000169604 |

| | | | |
|---|---|---|---|
| | | (already in table 1, other biological process) | |
| | **metastasis-associated fibroblasts (MAFs)** | CXCL12 (SDF1) | ENSG00000107562 |
| **Th1 response** | | TNF | ENSG00000232810 |
| **Macrophages /MDSC gene profile** | | NOS1 | ENSG00000089250 |
| | | IL8 (already in table 1, other biological process) | ENSG00000169429 |
| | | CD83 | ENSG00000112149 |
| | | CD86 | ENSG00000114013 |
| **T cell inhibition** | | IDO2 | ENSG00000188676 |
| | | IL7R | ENSG00000168685 |
| | | BTLA | ENSG00000186265 |
| | | NCAM1 | ENSG00000149294 |
| **General immunosuppression** | | BTLA (already in table 1, other biological process) | ENSG00000186265 |
| | | IDO2 (already in table 1, other biological process) | ENSG00000188676 |
| **Inflammatory** | | NOS2 (already in table 1, other biological process) | ENSG00000007171 |
| | | CD83 (already in table 1, other biological process) | ENSG00000112149 |
| **Natural killer cells** | | NCR1 | ENSG00000189430 |

| | | |
|---|---|---|
| | MICA | ENSG00000204520 |
| | MICB | ENSG00000204516 |
| | KLRK1 | ENSG00000213809 |
| | NCAM1 (already in table 1, other biological process) | ENSG00000149294 |
| **Dendritic cells** | ITGAM | ENSG00000169896 |

#### 2. Virtual H&E staining

The virtually stained hematoxylin and eosin (H&E) image, as shown in figure 1, is calculated from the DAPI-stained and the FITC-stained image using the method described in [3]:

$$\begin{aligned} R_{i,j} &= \exp\left[-k\left(a_R * D_{i,j} + b_R * F_{i,j}\right)\right] \\ G_{i,j} &= \exp\left[-k\left(a_G * D_{i,j} + b_G * F_{i,j}\right)\right] \\ B_{i,j} &= \exp\left[-k\left(a_B * D_{i,j} + b_B * F_{i,j}\right)\right] \end{aligned} \tag{1}$$

wherein $R_{i,j}$, $G_{i,j}$ and $B_{i,j}$ are the pixels of the red, the green and the blue channel of the H&E image, $D_{i,j}$ and $F_{i,j}$ are the pixels from the grey channel of the DAPI and the FITC image, respectively. The constants $k$, $a = (a_R, a_G, a_B)$ and $b = (b_R, b_G, b_B)$ are used for the colour and contrast adjustment of the virtually stained H&E image.

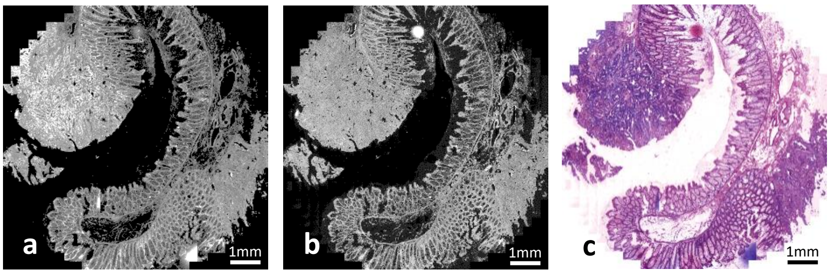


*Figure 1*: a) DAPI-stained image, b) FITC-stained image used for calculating and c) virtually stained H&E image of the tissue sample.

The parameters $a = (0.30, 1.00, 0.86)$ and $b = (0.54, 1.00, 0.05)$ are kept constant while the value of $k$ changes with each tissue sample so that a maximum value $p < 255$ is not exceeded in $R_{i,j}$, $G_{i,j}$ and $B_{i,j}$.

The virtually stained H&E images are then shown to a pathologist specialised on CRC, who classified the tissue in certainly neoplastic and non-neoplastic areas, i.e. by using the colours red and green respectively (see figure 2).

#### 3. Compartment building by morphology

For the computational tissue compartment (TC) building, areas with same colour are converted together into the same binary TC $C_{i,j}^{\text{tissue}}$, as shown in figure 2. Thereby, $C_{i,j}^{\text{tissue}}$ holds only the values 0 and 1 for all pixels

$$C_{i,j}^{\text{tissue}} = \begin{cases} 1 & \text{TC in pixel } (i,j) \\ 0 & \text{TC not in pixel } (i,j) \end{cases} \tag{2}$$

To take into account that particular areas $B_{i,j}$ of the image cannot be used for the analysis, e.g. due to too high autofluorescence of the tissue or lost tissue during hybridisation, these areas have to be excluded from all TC, i.e. by marking them with the colour blue as shown in figure 2:

$$C_{i,j}^{\text{analysis}} = C_{i,j}^{\text{tissue}} \setminus B_{i,j} \tag{3}$$

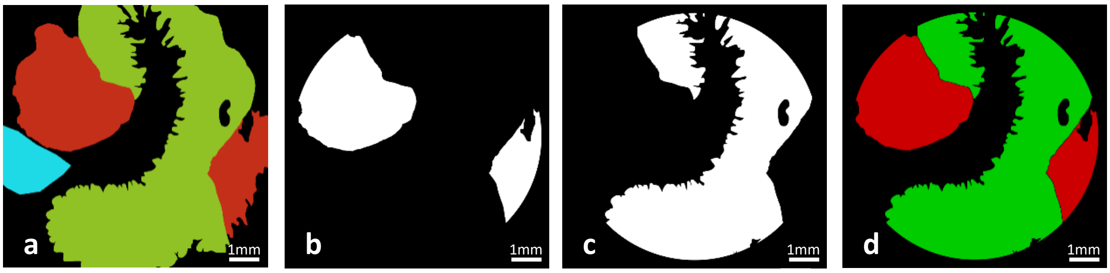


*Figure 2*: a) The tissue areas in the tissue sections as classified by a pathologist: red – neoplastic tissue, green – non-neoplastic tissue. The blue area marks a region that was excluded from the analysis due to high autofluorescence or lost tissue during hybridisation. The derived representative binary tissue compartment (TC) b) for the neoplastic and c) for the non-neoplastic tissue. d) The calculated TC combined in one image with the previously described colour coding.

### 4. Compartment building by gene expressions

In a pre-step, image analyses are performed on both microscope images of the pixel size $(I, J)$, on the *in situ* sequencing (ISS) images and on the DAPI-stained images:

The fractional image coordinates $\vec{x}_c$ for the nuclei $c$ centre positions of the cells $C$ and with 0<$\vec{x}_c$<1 are determined with the software tool CellProfiler™ through segmentation analysis. Further, the fractional image coordinates $\vec{x}_g$ for the positions of the ISS transcripts $g$ of gene $G$, e.g. HK1, are determined with the software MATLAB®.

Utilising $\vec{x}_g$, the distribution of all transcripts of gene $G$ in the tissue sample can be visualized through a density plot. For this purpose, each transcript is represented as disk-shaped element of radius $r$ with $\vec{x}_g$ as its centre' coordinate. A transcript's density $\rho_{i,j}^{g}$ at the pixel $(i, j)$ with the centre' coordinate $\vec{x}_{i,j}$ is then calculated by applying a uniform distribution kernel (see figure 3):

$$\rho_{i,j}^{g}(r) = \begin{cases} \frac{1}{2\pi r^2} & |\vec{x}_{i,j} - \vec{x}_g| < r \\ 0 & |\vec{x}_{i,j} - \vec{x}_g| \geq r \end{cases} \tag{4}$$

| | | | | | | |
|---|---|---|---|---|---|---|
| 0.0 | 0.0 | 0.0 | 1.0 | 0.0 | 0.0 | 0.0 |
| 0.0 | 1.0 | 1.0 | 1.0 | 1.0 | 1.0 | 0.0 |
| 1.0 | 1.0 | 1.0 | 1.0 | 1.0 | 1.0 | 1.0 |
| 1.0 | 1.0 | 1.0 | 1.0 | 1.0 | 1.0 | 1.0 |
| 1.0 | 1.0 | 1.0 | 1.0 | 1.0 | 1.0 | 1.0 |
| 0.0 | 1.0 | 1.0 | 1.0 | 1.0 | 1.0 | 0.0 |
| 0.0 | 0.0 | 0.0 | 1.0 | 0.0 | 0.0 | 0.0 |

*Figure 3*: Schematic example for the uniform kernel.

The density distribution $\rho_{i,j}^{G}$ for all transcripts of gene $G$ in the image is then given by the sum of all single transcript densities $\rho_{i,j}^{g}$:

$$\rho_{i,j}^{G}(r) = \sum_{\forall g \in G} \rho_{i,j}^{g}(r) \tag{5}$$

When the obtained density plots, as shown in in figure 4, are compared with the stained image of the tissue sample, some genes show a more or uniform density distribution in the whole tissue sample, e.g. FLT4 in neoplastic and non-neoplastic, while others show high densities only in certain tissue areas, e.g. MET predominately in neoplastic tissue.

For some genes, the high-density areas reveal a good correlation with the neoplastic tissue, as classified by a pathologist specialised on CRC in view of its morphological appearance. For example, the comparison of figure 2 and figure 4 shows that the distribution of BIK, EREG and MET can be associated with the neoplastic tissue.

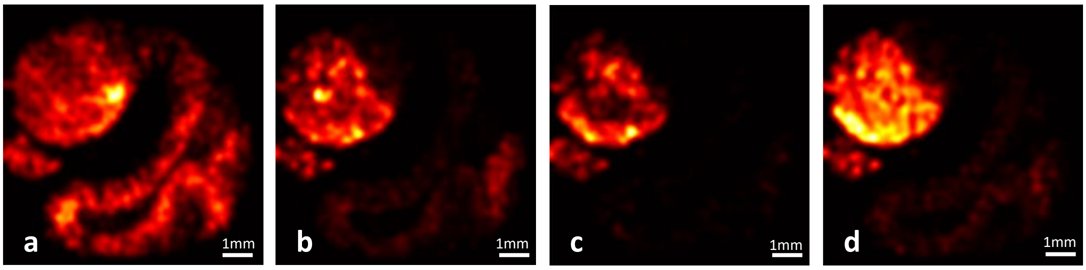


*Figure 4*: The density plots a) for FLT4, b) for BIK, c) for EREG and d) for MET. The areas with high density values (light red and yellow area) in b)-d) correlate with the areas of neoplastic

Based on this observation, a set of selected genes can be used for the automatic classification of the neoplastic area in tissue sections. Consequently, the remaining tissue represents then the non-neoplastic tissue.

In a first step, the genes with a high correlation are identified and combined to a gene set $S = \{G_1, G_2, \dots, G_n\}$. In a second step, the densities $\rho_{i,j}^{G}$ of the genes in $S$ are summed up to a total gene set density $\rho_{i,j}^{S}$, whereat $r$ is kept constant for all genes part of the set:

$$\rho_{i,j}^{S}(r) = \sum_{\forall G \in S} \rho_{i,j}^{G}(r) \tag{6}$$

Next, in order to convert the combined density plot into a TC $T_{i,j}$, a threshold is applied to the density value of each pixel $(i, j)$:

$$T_{i,j} = \begin{cases} 1 & \vartheta \leq \rho_{i,j}^{S}(r) \\ 0 & \vartheta > \rho_{i,j}^{S}(r) \end{cases} \tag{7}$$

Therein, the absolute threshold value $\vartheta$ is given as product of the relative threshold $\theta$ with the maximum value of $\rho_{i,j}^{G}$ in the whole image:

$$\vartheta = \theta \cdot \max_{I,J}[\rho_{i,j}^{S}] \quad \text{with } 0 \leq \theta \leq 1 \tag{8}$$

The threshold values used for the calculations are summarized in table .

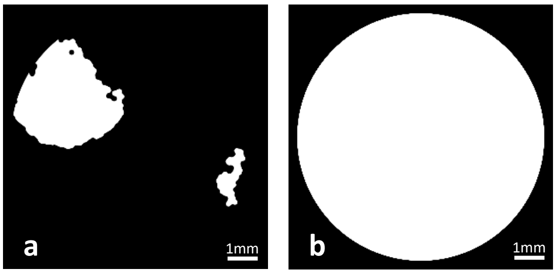


*Figure 4*: a) With the gene set *S = {BIK, CCND1, CD44, EREG, ITGAV, MET, MYBL2, S100A4},* referred to as tumour gene signature, calculated tissue compartment $T_{i,j}$ for the neoplastic tissue. b) Hybridisation area compartment $S_{i,j}$ defined through a disk of radius $R$ centered in the middle of image.

Disregarding the number of detected ISS transcripts and thus the resulting $\rho_{i,j}^{S}$, the values of the disk radius $r$ and of the relative threshold parameter $\theta$ determine the resolution of details of the generated TC. By increasing $r$, more and more disks merge, forming larger and connected areas. However, above a certain value, too many details of the tissue sample are lost. Similarly, a too high $\theta$ also causes a loss of details, while a too low value falsely depicts the TC by dispersed or displaced genes. With $\theta = 0.0$, all pixels $\rho_{i,j}^{S} > 0$ are part of the TC. The parameter values used for the calculations in this paper are summarized in table .

Further, to achieve an improvement of the binary TC, morphological transformations from the python library openCV [4] are used to fill up small gaps in the TC, i.e. by a closing operation followed by an immediate opening operation.

*Table 2*: Parameter values used for microscope images of size (7660px, 7700px).

| Tissue compartment | Disk radius *r* | Relative threshold $\theta$ |
|---|---|---|
| *neoplastic* | 180px | 0.10 |
| *composite cells* | 50px | 0.00 |
| *composite ISS genes* | 50px | 0.00 |

During the ISS hybridisation of the tissue, a secure-seal hybridization chamber (Sigma-Adrich) is mounted on every tissue sample. However, as the tissues are not well covered by the hybridisation solution near the chamber boundary, the number of detected transcripts is not representative in this area. Hence, a secure area $S_{i,j}$ with a certain indention from the secure seal boundary is defined where the number of detected transcripts is assumed to be representative for the tissue (see figure 4).

Thus, the secure area is defined as disk-shaped compartment:

$$S_{i,j}(R) = \begin{cases} 1 & \left|\vec{x}_{i,j} - \vec{C}\right| < R \\ 0 & \left|\vec{x}_{i,j} - \vec{C}\right| \geq R \end{cases} \tag{9}$$

where $\vec{C} = (I/2, J/2)$ and $R$ represent the centre of the image and the radius of the compartment, respectively.

The representative TC $C_{i,j}^{\text{tissue}}$ is then given through the intersection of $S_{i,j}$ with $T_{i,j}$:

$$C_{i,j}^{\text{tissue}} = T_{i,j} \cap S_{i,j}(R) \tag{10}$$

The non-neoplastic TC is obtained by building a compartment for the whole (composite) tissue sample $\mathrm{C}_{i,j}^{\text{composite}}$ and then subtracting the neoplastic one.

First, $\mathrm{C}_{i,j}^{\text{composite}}$ is calculated through intersection of the compartment $\mathrm{C}_{i,j}^{\text{ISS genes}}$ obtained from all detected ISS transcripts (of all genes) and the compartment $\mathrm{C}_{i,j}^{\text{cells}}$ derived from all cell nuclei positions, as shown in figure 5:

$$T_{i,j}^{\text{composite}} = T_{i,j}^{\text{cells}} \cap T_{i,j}^{\text{ISS genes}} \tag{11}$$

This approach reflects the fact that both tissue and ISS padlock reagents must be present for the correct detection of a gene. The parameters used for the calculations of the compartments are shown in table . Thereby, $\mathrm{T}_{i,j}^{\text{ISS genes}}$ as well as $\mathrm{T}_{i,j}^{\text{cells}}$ are calculated with the same procedure as described above for the neoplastic TC. However, due to the high number of observed ISS genes and cell nuclei, smaller values are used for $\mathrm{r}$ and $\theta$ to preserve the details of the tissue sample.

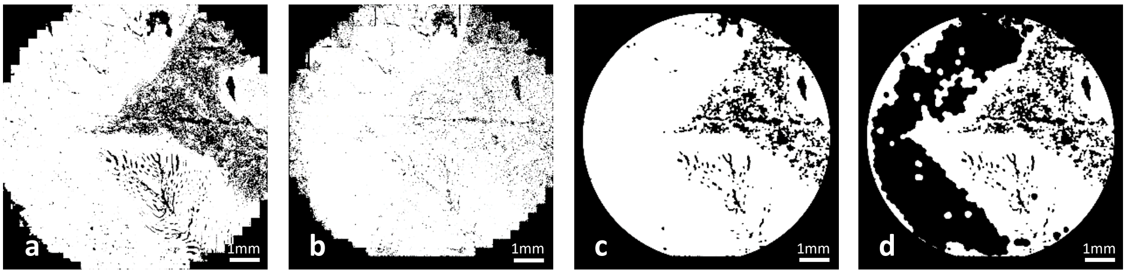


*Figure 5*: Binary tissue compartments (TC) a) for all cells $\mathrm{T}_{i,j}^{\text{cells}}$ and b) for all ISS genes $\mathrm{T}_{i,j}^{\text{ISS genes}}$. c) The composite TC $\mathrm{C}_{i,j}^{\text{composite}}$ and d) the calculated representative non-neoplastic TC.

The secure area of the composite TC $\mathrm{C}_{i,j}^{\text{composite}}$ is obtained by intersection of $\mathrm{T}_{i,j}^{\text{composite}}$ with the secure area compartment $S_{i,j}$, as described for the neoplastic tissue. In the next step, the representative non-neoplastic binary TC $\mathrm{C}_{i,j}^{\text{non-neoplastic}}$ is calculated through subtraction of the neoplastic TC from the composite TC:

$$C_{i,j}^{\text{non-neoplastic}} = C_{i,j}^{\text{composite}} \setminus C_{i,j}^{\text{neoplastic}} \tag{12}$$

To take into account that particular areas of the image $B_{i,j}$ cannot be used for the analysis, as described in the previous chapter, these areas have to be excluded from $C_{i,j}^{\text{non-neoplastic}}$ and $C_{i,j}^{\text{neoplastic}}$:

$$C_{i,j}^{\text{analysis}} = C_{i,j} \setminus B_{i,j} \tag{13}$$

Finally, figure *6* shows both obtained TC in one image whereby the same colour coding is used as in figure 2.

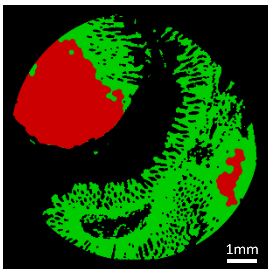


*Figure 6*: The calculated tissue compartments (TC) combined in one image: red – neoplastic TC and green – non-neoplastic TC.

#### 5. Gene counting in the compartments

The following function is used for the counting of transcripts or cells within a specific TC $C^{\text{tissue}}$ in which $\vec{x}_g$ is the fractional coordinate of the transcript's position in the image:

$$f(\vec{x}_g, C^{\text{tissue}}) = \begin{cases} 1 & \vec{x}_g \in C^{\text{tissue}} \\ 0 & \vec{x}_g \notin C^{\text{tissue}} \end{cases} \tag{14}$$

The number of transcripts of a certain gene $G$ per TC is then given through:

$$N_G^{\text{tissue}} = \sum_{\forall g \in G} f(\vec{x}_{g,} C^{\text{tissue}}) \tag{15}$$

Similarly, the number of cells per compartment is given by:

$$N_C^{\text{tissue}} = \sum_{\forall c \in C} f(\vec{x}_{c,} C^{\text{tissue}}) \tag{16}$$

The density per cell $\mathrm{D}_{\mathrm{G,C}}^{\text{tissue}}$ for the transcripts of gene $G$ is then given by:

$$\mathrm{D}_G^{\text{tissue}} = N_G^{\text{tissue}} / N_C^{\text{tissue}} \tag{17}$$

#### 6. Gene set selection

The overlap O between the morphological and the gene expression-based neoplastic TC, as shown in figure 7, is calculated for an image of the pixel size (I, J) via:

$$O = \frac{1}{A_{\text{morph}}^{\text{neoplastic}} A_{\text{gene}}^{\text{neoplastic}}} \sum_{\substack{\forall i \in I, \\ \forall j \in J}} \left( C_{i,j,\text{morph}}^{\text{neoplastic}} \cap C_{i,j,\text{gene}}^{\text{neoplastic}} \right)^2 \tag{18}$$

where $A_{\text{morph}}^{\text{neoplastic}}$ and $A_{\text{gene}}^{\text{neoplastic}}$ are the areas of the morphological and the gene expression-based neoplastic TC, respectively. The TC areas within are obtained by summing up all pixels of the TC:

$$A^{\text{tissue}} = \sum_{\substack{\forall i \in I, \\ \forall j \in J}} C_{i,j}^{\text{tissue}} \tag{19}$$

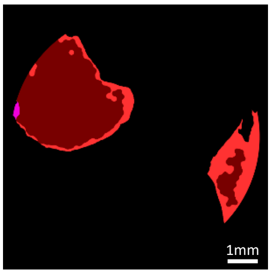


Figure 7: The overlap (dark red) between the neoplastic tissue compartment (TC): dark red – overlap, light red – morphological-based TC with no overlap and pink – gene expression-based TC with no overlap.

The composition of the gene set $S$ is alternated and the overlaps are calculated for all $N$ patient samples. The mean overlap used is then used for the rating of $S$:

$$O_{\text{mean}} = \sqrt[N]{O_1 O_2 \ldots O_N} \tag{20}$$

Finally, the gene set consisting of the 8 genes *S = {BIK, CCND1, CD44, EREG, ITGAV, MET, MYBL2, S100A4}* and referred to as tumour gene signature, achieved the highest $O_{\text{mean}}$ value is therefore selected for statistical testing.

### 7. Statistical testing

A two-tailed paired t-test is used for the statistical significance testing of the transcript distribution in the neoplastic $\mathrm{D}_G^{\mathrm{neoplastic}}$ and the non-neoplastic $\mathrm{D}_G^{\mathrm{non-neoplastic}}$ tissues of the $\mathrm{N}$ patients:

$$\begin{aligned} H_0&: \mu_d = 0 \\ H_1&: \mu_d \neq 0 \end{aligned} \tag{21}$$

with $d_n = \mathrm{D}_G^{\mathrm{neoplastic}} - \mathrm{D}_G^{\mathrm{non-neoplastic}}$ for $\alpha/2 = 0.05$.

Further, a two-tailed independent t-test is applied for statistical significance testing of the transcript distribution in the neoplastic tissue of the $N/2$ patients in the relapse group with the mean $\mu_{\mathrm{relapse}} = \overline{\mathrm{D}}_{G,\mathrm{relapse}}^{\mathrm{neoplastic}}$ and the non-relapse group with the mean $\mu_{\mathrm{no-relapse}} = \overline{\mathrm{D}}_{G,\mathrm{no-relapse}}^{\mathrm{neoplastic}}$, yielding:

$$\begin{aligned} H_0&: \mu_{\mathrm{relapse}} = \mu_{\mathrm{no-relapse}} \\ H_1&: \mu_{\mathrm{relapse}} \neq \mu_{\mathrm{no-relapse}} \end{aligned} \tag{22}$$

for $\alpha/2 = 0.05$.

The statistical testing is done for both the morphological as well as for the by gene expression-based TC.

*Table 3:* The resulting p-values for the statistical testing of relapse and non-relapse patients with the neoplastic tissue compartments.

| genes | p-value | mean relapse patients | mean non-relapse patients |
|---|---|---|---|
| ADGRA2 | 0.4380 | 0.0017 | 0.0013 |
| ALDH1A1 | 0.9271 | 0.0021 | 0.0019 |
| ANGPT1 | 0.9514 | 0.0010 | 0.0010 |
| ANGPT2 | 0.2625 | 0.0104 | 0.0040 |
| ANPEP | 0.5374 | 0.0021 | 0.0034 |
| ANTXR1 | 0.5995 | 0.0764 | 0.0673 |
| ATG5 | 0.4726 | 0.0041 | 0.0029 |
| AURKA | 0.7366 | 0.0092 | 0.0083 |
| BAK1 | 0.9219 | 0.0087 | 0.0084 |
| BBC3 | 0.8933 | 0.0893 | 0.0937 |
| BCL2L1 | 0.1268 | 0.0601 | 0.0360 |
| BCL2L11 | 0.7712 | 0.0039 | 0.0041 |
| BECN1 | 0.6856 | 0.0043 | 0.0036 |
| BEST4 | 0.3816 | 0.0058 | 0.0046 |
| BGN | 0.5344 | 0.0093 | 0.0065 |
| BID | 0.2140 | 0.0429 | 0.0200 |
| BIK | 0.4501 | 0.0260 | 0.0188 |
| BMI1 | 0.3415 | 0.0084 | 0.0020 |
| BOP1 | 0.8287 | 0.0327 | 0.0366 |
| BTG2 | 0.1062 | 0.0519 | 0.0287 |
| BTLA | 0.4055 | 0.0005 | 0.0006 |
| CA1 | 0.3870 | 0.0022 | 0.0017 |
| CA2 | 0.5645 | 0.0012 | 0.0017 |
| CASP3 | 0.0700 | 0.0100 | 0.0057 |
| CASP7 | 0.0682 | 0.0609 | 0.0210 |
| CASP8 | 0.5583 | 0.0034 | 0.0024 |
| CASP9 | 0.2590 | 0.0041 | 0.0028 |
| CAT | 0.8154 | 0.0054 | 0.0050 |
| CCND1 | 0.8206 | 0.0243 | 0.0207 |
| CCNE1 | 0.5620 | 0.0411 | 0.0279 |
| CD248 | 0.4689 | 0.0030 | 0.0037 |
| CD44 | 0.9015 | 0.0186 | 0.0196 |
| CD80 | 0.6438 | 0.0001 | 0.0001 |
| CD83 | 0.5719 | 0.0016 | 0.0018 |
| CD86 | 0.0738 | 0.0011 | 0.0006 |
| CDH5 | 0.2937 | 0.0026 | 0.0019 |
| CFLAR | 0.2040 | 0.0032 | 0.0008 |
| CHGA | 0.1433 | 0.0076 | 0.0038 |
| CNTD2 | 0.5664 | 0.0091 | 0.0124 |
| COL1A2 | 0.6891 | 0.1772 | 0.1424 |
| CSPG4 | 0.3571 | 0.0077 | 0.0057 |
| CXCL1 | 0.2984 | 0.0069 | 0.0042 |

| | | | |
|---|---|---|---|
| CXCL12 | 0.8857 | 0.0076 | 0.0073 |
| CXCL8 | 0.4133 | 0.0058 | 0.0039 |
| DCLK1 | 0.2134 | 0.0012 | 0.0008 |
| DCN | 0.7197 | 0.0016 | 0.0018 |
| E2F1 | 0.8938 | 0.0094 | 0.0088 |
| EGFR | 0.3622 | 0.0165 | 0.0109 |
| EGLN3 | 0.2134 | 0.0066 | 0.0125 |
| ENAH | 0.2609 | 0.0263 | 0.0400 |
| ENG | 0.6169 | 0.0630 | 0.0534 |
| ENTPD1 | 0.4883 | 0.0013 | 0.0049 |
| EPHB2 | 0.1347 | 0.0030 | 0.0017 |
| EREG | 0.2798 | 0.0085 | 0.0037 |
| EXOSC5 | 0.5477 | 0.0307 | 0.0241 |
| F8 | 0.5569 | 0.0041 | 0.0027 |
| FABP1 | 0.1063 | 0.3312 | 0.1101 |
| FABP4 | 0.2537 | 0.0015 | 0.0006 |
| FAM3C | 0.5082 | 0.0042 | 0.0033 |
| FGFR2 | 0.0137 | 0.0049 | 0.0031 |
| FLT4 | 0.3144 | 0.0252 | 0.0179 |
| FN1 | 0.2072 | 0.0830 | 0.0256 |
| FOXC2 | 0.1784 | 0.0009 | 0.0006 |
| FRK | 0.7101 | 0.0015 | 0.0014 |
| FSTL1 | 0.7359 | 0.0068 | 0.0063 |
| GADD45B | 0.3688 | 0.0076 | 0.0062 |
| GAST | 0.0856 | 0.0046 | 0.0032 |
| GFI1B | 0.4393 | 0.0023 | 0.0014 |
| GIP | 0.1603 | 0.0011 | 0.0007 |
| GLS | 0.5713 | 0.0131 | 0.0104 |
| GLUD1 | 0.6640 | 0.0028 | 0.0032 |
| GPX1 | 0.6236 | 0.0280 | 0.0229 |
| GRB7 | 0.2775 | 0.0115 | 0.0026 |
| GSR | 0.6764 | 0.0213 | 0.0176 |
| GUCA2B | 0.2908 | 0.0024 | 0.0012 |
| HIF1A | 0.7241 | 0.0043 | 0.0048 |
| HIF3A | 0.1704 | 0.0037 | 0.0020 |
| HK1 | 0.4892 | 0.0145 | 0.0115 |
| HK2 | 0.5087 | 0.0157 | 0.0117 |
| HMGB1 | 0.3301 | 0.0080 | 0.0023 |
| HYOU1 | 0.4569 | 0.0457 | 0.0344 |
| ICAM1 | 0.7122 | 0.0051 | 0.0044 |
| IDO2 | 0.1770 | 0.0009 | 0.0004 |
| IGFBP2 | 0.7922 | 0.0483 | 0.0441 |
| IL11 | 0.3020 | 0.0034 | 0.0027 |
| IL1B | 0.1749 | 0.0265 | 0.0187 |
| IL6 | 0.3527 | 0.0019 | 0.0009 |
| IL7R | 0.4805 | 0.0017 | 0.0015 |
| INHBA | 0.9982 | 0.0038 | 0.0038 |

| | | | |
|---|---|---|---|
| ITGAM | 0.8851 | 0.0022 | 0.0021 |
| ITGAV | 0.6371 | 0.0158 | 0.0126 |
| ITGAX | 0.3278 | 0.0010 | 0.0005 |
| KDR | 0.7293 | 0.0027 | 0.0024 |
| KIT | 0.4801 | 0.0010 | 0.0005 |
| KLF4 | 0.1445 | 0.2326 | 0.0777 |
| KLRK1 | 0.8630 | 0.0007 | 0.0007 |
| L1CAM | 0.1590 | 0.0040 | 0.0031 |
| LAMC2 | 0.5822 | 0.0051 | 0.0078 |
| LDHA | 0.1784 | 0.0052 | 0.0081 |
| LGR5 | 0.7745 | 0.0043 | 0.0038 |
| LIF | 0.3691 | 0.0055 | 0.0034 |
| MAP1LC3A | 0.6020 | 0.0214 | 0.0168 |
| MCL1 | 0.3200 | 0.0178 | 0.0114 |
| MCM2 | 0.4943 | 0.0118 | 0.0084 |
| MET | 0.4367 | 0.0327 | 0.0499 |
| MICA | 0.4789 | 0.0023 | 0.0030 |
| MICB | 0.2659 | 0.0157 | 0.0062 |
| MIEN1 | 0.1606 | 0.0099 | 0.0029 |
| MLKL | 0.9694 | 0.0375 | 0.0370 |
| MLN | 0.2325 | 0.0053 | 0.0019 |
| MMP11 | 0.0415 | 0.0492 | 0.0164 |
| MMP2 | 0.8319 | 0.0019 | 0.0021 |
| MMP7 | 0.8832 | 0.0061 | 0.0068 |
| MPL | 0.1839 | 0.0010 | 0.0006 |
| MUC2 | 0.2674 | 0.0124 | 0.0080 |
| MUC5AC | 0.2946 | 0.0008 | 0.0005 |
| MYBL2 | 0.4202 | 0.0091 | 0.0057 |
| NANOG | 0.2675 | 0.0042 | 0.0024 |
| NCAM1 | 0.6225 | 0.0117 | 0.0106 |
| NCR1 | 0.7844 | 0.0010 | 0.0010 |
| NOS1 | 0.4461 | 0.0129 | 0.0102 |
| NOS2 | 0.6146 | 0.0076 | 0.0109 |
| NT5E | 0.8994 | 0.0027 | 0.0025 |
| NTS | 0.3922 | 0.0013 | 0.0008 |
| OLFM4 | 0.6367 | 0.0911 | 0.1395 |
| OSER1 | 0.4527 | 0.0102 | 0.0069 |
| OTOP2 | 0.0043 | 0.0026 | 0.0019 |
| PCNA | 0.5341 | 0.0125 | 0.0077 |
| PDGFA | 0.1237 | 0.0302 | 0.0157 |
| PDK1 | 0.4882 | 0.0024 | 0.0032 |
| PKM | 0.7688 | 0.0922 | 0.0817 |
| PLXDC1 | 0.5181 | 0.0062 | 0.0051 |
| POU2F3 | 0.5070 | 0.1033 | 0.0636 |
| POU5F1 | 0.5592 | 0.5020 | 0.3878 |
| PPIF | 0.3442 | 0.0056 | 0.0035 |
| PRDX2 | 0.7580 | 0.1062 | 0.0872 |

| | | | |
|---|---|---|---|
| PRF1 | 0.2779 | 0.0005 | 0.0011 |
| PROM1 | 0.0792 | 0.0086 | 0.0026 |
| PYY | 0.3809 | 0.0027 | 0.0019 |
| RIPK1 | 0.7866 | 0.0047 | 0.0051 |
| RIPK3 | 0.2560 | 0.0153 | 0.0115 |
| RPS6KB1 | 0.5633 | 0.0004 | 0.0008 |
| S100A4 | 0.0645 | 0.0308 | 0.0118 |
| SAAL1 | 0.3395 | 0.0018 | 0.0011 |
| SALL4 | 0.6568 | 0.0019 | 0.0023 |
| SCAI | 0.1913 | 0.0047 | 0.0026 |
| SLC26A3 | 0.9402 | 0.1045 | 0.1083 |
| SLC2A1 | 0.3493 | 0.0006 | 0.0004 |
| SOD1 | 0.9646 | 0.0983 | 0.0971 |
| SOX2 | 0.8878 | 0.0138 | 0.0131 |
| SPDEF | 0.2998 | 0.0039 | 0.0023 |
| SST | 0.5133 | 0.0011 | 0.0007 |
| TAGLN | 0.1000 | 0.0039 | 0.0015 |
| TBXT | 0.4052 | 0.0027 | 0.0012 |
| TEK | 0.9708 | 0.0005 | 0.0005 |
| TFF1 | 0.5438 | 0.0092 | 0.0134 |
| TFF3 | 0.4431 | 0.0703 | 0.0396 |
| TGFA | 0.1879 | 0.0057 | 0.0034 |
| TGFB3 | 0.9421 | 0.0032 | 0.0033 |
| THY1 | 0.6415 | 0.0036 | 0.0030 |
| TIAM1 | 0.2424 | 0.0009 | 0.0006 |
| TIE1 | 0.2877 | 0.0013 | 0.0010 |
| TIMP1 | 0.9941 | 0.0257 | 0.0256 |
| TJP1 | 0.4679 | 0.0133 | 0.0079 |
| TNC | 0.8990 | 0.0073 | 0.0068 |
| TNF | 0.5561 | 0.0010 | 0.0006 |
| TNFRSF10A | 0.4299 | 0.0071 | 0.0035 |
| TNFRSF10B | 0.3869 | 0.0028 | 0.0041 |
| TNFSF10 | 0.8549 | 0.0077 | 0.0071 |
| TRPM5 | 0.3110 | 0.0058 | 0.0025 |
| TWIST1 | 0.5346 | 0.0256 | 0.0212 |
| TXNL1 | 0.3908 | 0.0069 | 0.0049 |
| URGCP | 0.5626 | 0.0002 | 0.0004 |
| VCAM1 | 0.9583 | 0.0007 | 0.0007 |
| ZEB1 | 0.2654 | 0.0014 | 0.0007 |
| ZGLP1 | 0.3593 | 0.0026 | 0.0046 |

### 8. Literature citations


1. Uhlén M, Fagerberg L, Hallström BM, Lindskog C, Oksvold P, Mardinoglu A, Sivertsson Å, Kampf C, Sjöstedt E, Asplund A, Olsson I, Edlund K, Lundberg E, Navani S, Szigyarto CA-K, Odeberg J, Djureinovic D, Takanen JO, Hober S, et al.: **Proteomics. Tissue-based map of the human proteome.** *Science* 2015, **347**:1260419.

2. Human Protein Atlas proteinatlas.org

3. Giacomelli MG, Husvogt L, Vardeh H, Faulkner-Jones BE, Hornegger J, Connolly JL, Fujimoto JG: **Virtual Hematoxylin and Eosin Transillumination Microscopy Using Epi-Fluorescence Imaging**. *PLoS One* 2016; 11:e0159337

4. Bradski G: **The OpenCV Library**. *Dr Dobb's Journal of Software Tools* 2000
